# Inter-Instrument Quantification of Fluorescence Read-out Signal Using DNA Origami Calibrator Beads

**DOI:** 10.1101/2025.11.13.683362

**Authors:** A.B. Bohn, C.C. Petersen, J. Wood, F.S. Pedersen, D. Selnihhin

## Abstract

Comparing experimental results across different instruments and laboratories is challenging due to variations in instrument sensitivity and resolution, particularly for analyzing nano-sized and dimly fluorescing particles like viruses and extracellular vesicles. To address this challenge, we conducted a study using fluorescent DNA calibrator beads to test, calibrate, and compare the sensitivity of five different flow cytometers.

Our approach involves DNA beads produced through a bottom-up DNA origami method. These beads contain a defined number of fluorophores (0-220) and exhibit no autofluorescence.

Fluorophores are precisely conjugated onto the beads in a controlled array, minimizing dye interactions. This allows triggering on one fluorophore and calibration based on another, facilitating precise and accurate calibration in absolute fluorophore numbers.

We tested five flow cytometers—FACSAria III, NovoCyte 3000YBG, NovoCyte Quanteon 4500, LSRFortessa, and CytoFlex S—using Cy5 and FAM DNA calibrator beads. We investigated various instrument parameters, including count rate, trigger and calibration gain, gain amplification linearity, and fluorescence trigger sensitivity. Using fluorescent triggering, we demonstrated the capability to detect dimly fluorescent particles, enabling instrument calibration and fluorescence sensitivity comparison using DNA calibrators.

In conclusion, our study offers a robust method for addressing instrument fluorescence channel resolution in absolute fluorophore numbers in the analysis of nano-sized and dimly fluorescent particles.

## Introduction

A major contributor to the complexity of comparing experimental results between instruments and laboratories is differences in instrument sensitivity and resolution. Knowing your instrument’s fluorescence channel resolution in absolute fluorophore numbers and comparing this information between instruments is of great value for many laboratories, especially for those analyzing small particles and/or dimly fluorescent cells/particles. This information is relevant not only when choosing which instrument/detector to use for a given analysis but also when reporting data from e.g. small particle analysis like viruses or extracellular vesicles. In this study, we use fluorescence calibrator DNA beads [1], with up to 220 fluorophores per particle, as well as Spherotech Molecules of Equivalent Soluble Fluorochrome (MESF) beads, with 1,322 – 787,489 MESF per particle, to determine and compare the sensitivity of five different flow cytometers.

Commercial beads are available for this purpose and can be used for as well the calibration as the estimation of instrument resolution (Q value) and sensitivity (B value) [2]. The B value is dependent on a combination of optic and electronic noise, due to experimental conditions, whereas the Q value depends on a combination of the number of photoelectrons generated per molecule of excited fluorochrome [2, 3] and the efficacy of the instrument to detect these photons, convert them to photoelectrons, and generate an electronic signal [3, 4]. The lower the B value, the better the detection potential of a positive signal, and the higher the Q value, the better the separation between populations. The Q and B values can, therefore, provide information about the performance of each detector on an instrument and can be used to compare instrument-to-instrument variation or as a quality assurance [4].

However, there are a few limitations to the commercially available beads. They are made of plastic materials, which introduces inaccuracies due to their autofluorescence and thereby high background [5]. This limits the quantification of a low number of fluorophores present on e.g. viruses or extracellular vesicles. The number of fluorophores on these plastic beads is reported as molecules of equivalent soluble fluorochrome (MESF) units and thus, this does not give an absolute number of fluorophores per bead but is an estimation of the number of conjugated fluorophores obtained by comparing bead fluorescence to the number of free fluorophores in solution [6–8]. Given that fluorophores in solution may have different molar absorption from fluorophores conjugated to beads [6, 7], they may not provide an accurate number of fluorochromes per particle. Furthermore, the fluorophores in these internally dye-stained plastic beads are not site-directed and have variable distances between fluorophores that can lead to dye-dye interactions resulting in homo-FRET (Förster resonance energy transfer), FRET, and quenching [9].

Moreover, these commercially available beads were originally developed for cell analysis and the lowest number of fluorophores are, to our knowledge, around hundreds to thousands of fluorophores probably highly exceeding the expected number of fluorochromes present on small particles. Therefore, reporting instrument resolution in small particle analysis, based on currently commercially available beads, relies on an extrapolation which may introduce inaccuracies in terms of background (sensitivity) and the number of fluorochromes present on small particles [10].

In a recently published paper [1], we presented the development and validation of precise and accurate fluorescence DNA calibrator beads applicable for the analysis of small particles, such as viruses, using flow cytometry. These DNA beads are produced by a bottom-up approach using a DNA origami method [11–13] that allows the synthesis of beads containing a precise even number, between 0 and 220, of fluorochromes per DNA calibrator bead. The fluorophores are conjugated onto DNA beads in a highly regular array with distances between fluorophores that prevent dye-dye interaction [9]. These DNA beads provide means for the translation of fluorescence intensity signal into absolute fluorochrome numbers. One important advantage of these DNA beads is that they exhibit essentially no autofluorescence. Moreover, two different fluorophores arranged in modular fluorophore arrays can be attached to the beads enabling us to trigger on one fluorescence parameter and perform fluorescence detection and calibration based on the other fluorophore. Therefore, a calibration curve including DNA beads with zero fluorochromes in the calibration module allows for a direct and precise determination of the background fluorescence/detection limit for a given flow cytometric detector due to a precise intercept with the y-axis.

In this study, the DNA beads were used to calibrate 5 different instruments (FACSAria III, NovoCyte 3000YBG, NovoCyte Quanteon 4500, LSRFortessa, and CytoFlex S). We used Cy5 and 5(6)-carboxyfluorescein (FAM) DNA calibrator beads [1] to test several parameters of the instruments, such as the highest count rate, optimal trigger and calibration gain, detector gain amplification linearity, and fluorescence trigger sensitivity. As expected, only some of the instruments were able to detect single DNA beads. The calibration of the instruments that could detect single DNA beads was compared to calibration based on two sets of commercially available beads (APC and FITC MESF beads from Spherotech).

## Results

Initially, the maximum count rate of each flow cytometer (FCM) was determined, where singular events correspond to the detection of single DNA beads (Figure S1). Two types of beads, labeled with either 10 or 100 FAM dyes for calibration and both labeled with 100 Cy5 dyes for triggering, were used for this experiment. A bead with 100 Cy5 and 0 FAM dyes was used to define the background fluorescence in the FAM detector of each FCM shown as a black dashed line in Figure S1. Event detection was triggered by Cy5 fluorescence, and the FAM-H median fluorescence intensity (MFI) was recorded and plotted against particle dilution. To determine the maximum count rate avoiding swarming we performed dilution controls. Single particle detection is obtained when the MFI values remain stable, and the count rate decreases in correlation with dilution. Swarming was evident for the LSRFortessa at the higher bead concentrations where several bead particles were detected as one event leading to higher median fluorescence intensity (MFI) per detected event (Figure S1 **a** and **c**). Dilution led to a lowering of the MFI value and increased count rate (Figure S1 **b** and **d**). A 16-times dilution of the stock bead solution resulted in the stabilization of MFI and a proportional decrease in count rate because of the detection of one bead per triggered event (Figure S1 **c** and **d**). The maximum count rate for the LSRFortessa corresponded to an average count rate of 6,500 events/s. A higher count rate may indicate that conditions for swarming could be present. This maximum count rate is similar to what we have earlier defined as the maximum count rate speed for the Cytoflex [1].

Although beads with 10 or 100 FAM dyes for the calibration show similar results, beads with 100 FAM dyes are more sensitive in this type of experiment because of the higher MFI and better separation from the background (Figure S1a and S1c). The Quanteon showed stable MFI and a linear decrease in count rate over the entire dilution interval, without any significant swarming (Figure S1 e-h), achieving a maximum count rate of more than 15,000 events/s. The maximum recorded count rate of the NovoCyte (Figure S1 i-l) and the FACSAria III (Figure S1 m-p) was just above 2,150 events/s and 8,400 events/s, respectively, without showing any significant swarming. To compare the instruments objectively, all FCMs should be compared at their most sensitive setting. Our next objective was, therefore, to determine the most sensitive gain for both triggering and detection for each of the FCMs. We varied the trigger gain while keeping the calibration gain constant. The most sensitive trigger gain resulted in the highest count rate, allowing the detection of most beads in the solution (Figure S2 and S3).

The optimal FAM trigger gain for the LSRFortessa was 630 where the highest count rate was achieved (Figure S2 **a** and **b**), for the Quanteon it was 800 (Figure S2 **c** and **d**), 575 for the Novocyte (Figure S2 **e** and **f**), 700 for the FACSAria III (Figure S2 **g** and **h**), and 1,500 for the Cytoflex (Figure S2 **i** and **j**). It is important to note that total background fluorescence determined by DNA beads with 0 Cy5 dyes stayed at the same level independent of the FAM trigger gain and equal to MFI of the background set by buffer alone (Figure S2 **b, d, f, h**, and **j**). Similarly, the most sensitive calibration gain was identified by keeping the trigger gain constant at the optimal gain, determined above, and only varying the calibration gain. The MFI of the calibration fluorophore was plotted against calibration gain and the most sensitive detector gain was defined as the gain providing the highest signal-to-background ratio.

Increasing the Cy5 trigger gain on the LSRFortessa resulted in a decreasing count rate and increasing MFI of detected events (Figure S3 **a** and **b**). Increasing the trigger gain led to the detection of fewer particles with higher fluorescence per particle, thus only the detection of larger oligomers that are also present to a smaller degree (Figure S10 and [14]). The highest count rate, and thus the optimal Cy5 trigger gain, was 560 for the LSRFortessa, 800 for the Quanteon (Figure S3 **c** and **d**), 600 for the Novocyte (Figure S3 **e** and **f**), 800 for the FACSAria III (Figure S3 **g** and **h**), and 400 for the Cytoflex (Figure S3 **i** and **j**). The total background fluorescence, determined by DNA beads with 0 FAM dyes, was independent of the Cy5 trigger gain and equal to the MFI of the background set by buffer alone (Figure S3 **b, d, f, h**, and **j**).

To assess the linearity of detector gain for the different FCMs, we used the MFI in the detector channel of beads with the trigger fluorophores and zero detector fluorophores as the background signal (Figure S4 and S5). This is possible because DNA has a maximum absorbance at 260 nm and a very low quantum yield and extinction coefficient [15] compared to fluorophores and, thus, does not have any significant fluorescence at FAM (Figure S6) and Cy5 (Figure S7) emission wavelengths. Therefore, the emission recorded from the DNA beads without any fluorophore in the calibration module represents a true background of the FCM.

Higher FAM gain leads to higher FAM median fluorescence intensity for all tested FCMs (Figure S4). On the Cytoflex, an increase in FAM gain resulted in a linear amplification of fluorescence intensity, as evident from the trendline (Figure S4, green dashed line) for 100 FAM bead data and a corresponding R-squared value. The R-squared value is similar for the other DNA beads tested.

The signal-to-background ratio was calculated to determine the gain providing the highest resolution where beads with 0 FAM were used to determine the total background. The highest resolution was achieved at a gain of 1000 for the LSRFortessa and the FACSAria III, at a gain of 800 for the Quanteon and the Novocyte, and a gain of 2000 for the Cytoflex. In our experiments, the Cytoflex was shown to have absolute linear gain amplification with R^2^ > 0.99, the Quanteon with R^2^ > 0.94, while the other tested FCMs had R^2^ < 0.90 (Figure S4 and S5). Absolute linear gain amplification allows the recalculation of MFI values for other gain settings.

Similarly, we determined the optimal Cy5 detector gain (Figure S5). The higher Cy5 gain led to higher Cy5 median fluorescence intensity for all tested FCMs. On the Cytoflex, an increase of Cy5 gain resulted in linear amplification of Cy5 fluorescence intensity, as evident from the trend line (green dashed line) for 110 Cy5 bead data and a corresponding R-squared value that is representative also for other DNA beads. The signal-to-background ratio was calculated to determine the highest resolution gain. Beads with 0 Cy5 were used to determine the total background. The highest resolution was achieved at a gain of 1,000 for the LSRFortessa, the Quanteon, and the FACSAria III, at a gain of 800 for the Novocyte, and a gain 2,500 for the Cytoflex.

Next, we determined how many fluorophores were needed in the trigger module for an FCM setup to be able to detect single beads. For this purpose, we assembled DNA beads with 70 Cy5 fluorophores in the calibration module and varying numbers (10-140) of FAM fluorophores in the trigger module (Figure S8). At a concentration of 215 pM, the beads were analyzed on different FCMs using the determined optimal trigger/calibration gain settings (Figure S2-S5). To prevent swarming, the beads were further diluted, if necessary, to not exceed the maximum count rate determined previously (Figure S1). All beads were run at optimal detector gain, as determined earlier and shown in Figure S2 with a maximum count rate not exceeding 4,500 events/s (2,000 events/s for Novocyte) to prevent swarming. Beads with 100 FAM fluorophores in the trigger module and no Cy5 fluorophores were used to determine the background in the Cy5 detector.

Analysis of densitometric data, obtained from electrophoretic analysis of the assembled DNA beads, revealed that 74% of the particles were dimers as designed (Figure S10). The remaining particles consist of monomers (12%), larger oligomers (14%) of trimers/tetramers, and probably even larger oligomers in smaller amounts that are not detectable on the gel. At low fluorophore trigger numbers, only the large oligomers are detected, resulting in high Cy5-H MFI values of the detected events (Figure S8). The detection of beads with 10 FAM fluorophores in the trigger module was poor: ≤1%. Only rare bead oligomers (Figure S10) were detected when particle detection was triggered by these 10 FAM fluorophore arrays. Therefore, the background contributed largely to event count, thus decreasing MFI values. Increasing the number of the trigger fluorophores on the DNA beads results in the detection of smaller oligomers and homodimers, which leads to a decrease in Cy5-H MFI values and an increase in the number of detected particles in the DNA bead solution (Figure S8a). Detected concentrations of each sample were calculated using the count rate and FCM flow rate. The LSRFortessa was able to detect 1.3% of all beads when DNA beads with 140 FAM in the trigger module were used. This number was 44.1% for the Quanteon, 7.3% for the Novocyte, 4.9% for the FACSAria III, and 80.1% for the Cytoflex.

A similar experiment was performed using DNA beads with varying numbers (10-160) of Cy5 fluorophores in the trigger module and 60 FAM molecules in the calibration module (Figure S9). The LSRFortessa was able to detect 39.3%, the Quanteon 49.8%, Novocyte 2.8%, FACSAria III 71.2%, and Cytoflex 11.5% of all DNA beads labelled with 160 Cy5 fluorophores in the trigger module and 60 FAM molecules in the calibration module.

Only the Cytoflex FCM was able to detect more than half of all beads based on 140 FAM fluorophores as a trigger and the FACSAria III was able to detect more than half of all beads when triggered by 160 Cy5 fluorophores. To investigate the sensitivity of these two FCMs further, we assembled beads with 160 Cy5 fluorophores in the trigger module and varying numbers of FAM fluorophores in the calibration module: 0, 14, 30, 44, and 60 FAM. Similarly, beads with 150 FAM in the trigger module were labeled with an increasing number of Cy5 fluorophores: 0, 14, 30, 60, and 70 Cy5. Furthermore, we included the Quanteon that performed similarly well in the detection of DNA beads for Cy5 and FAM trigger with a detection level of 44-50% of all beads. We also included commercially available FITC and APC calibration particles from Spherotech to compare these to the DNA beads (Figure 1). Although FITC and FAM are not the same fluorophores, these are both derivatives of fluorescein and have almost identical physical properties: very similar excitation/emission profiles, extinction coefficient, and quantum yield. APC and Cy5 are, on the other hand, two very different fluorophores, and the only similarity is that they are detected in the same wavelength area. The aim was to assess if calibration based on APC beads results in the same y-axis intersection as calibration based on Cy5 DNA beads since DNA beads lack any significant autofluorescence and thus provide real FCM background fluorescence.

**Figure 1.**
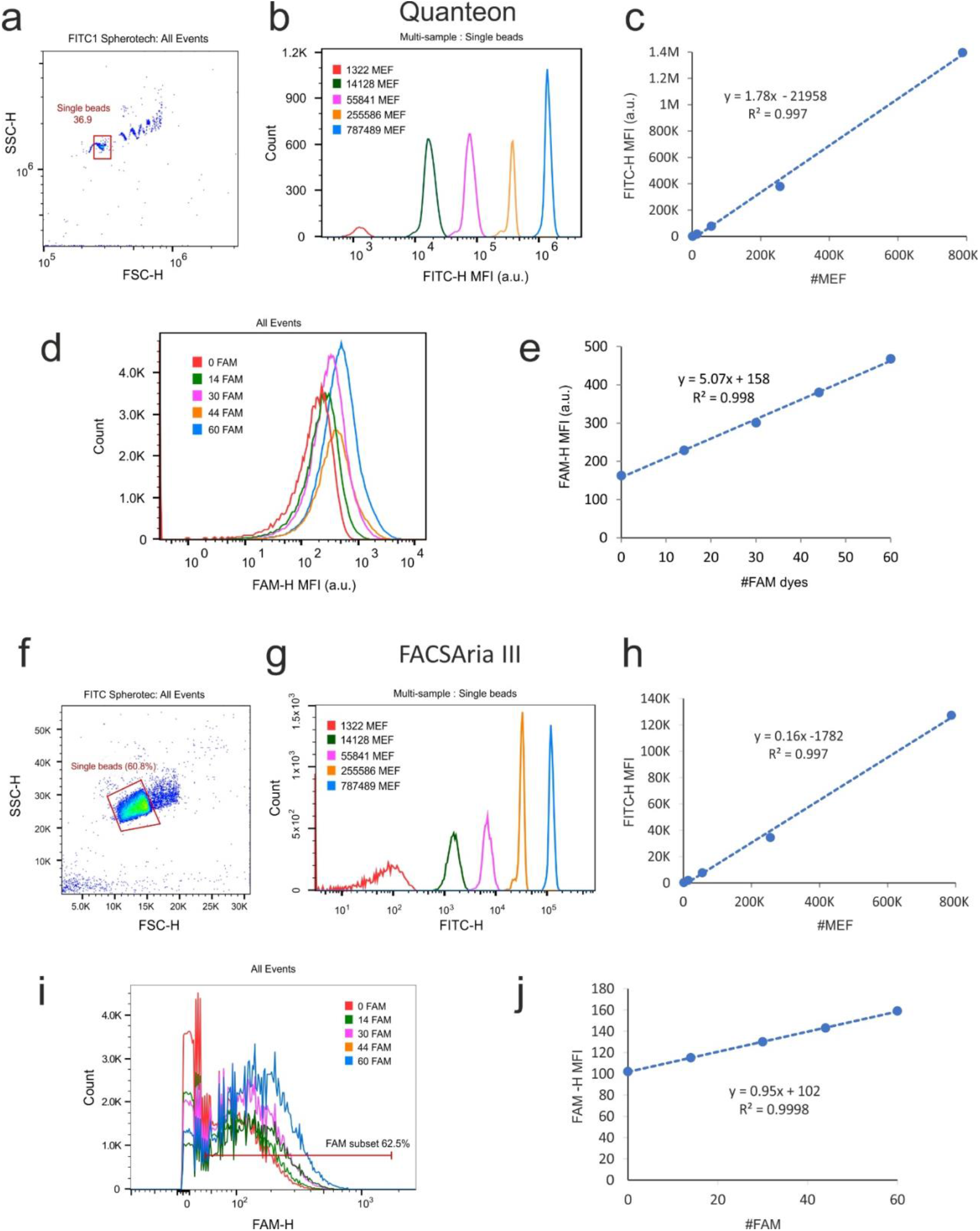
Calibration of the Quanteon (**a-e**) and the FACSAria III (**f-j**) using FITC Spherotech and FAM DNA calibration beads. **a and f**. Spherotech single FITC beads were gated using Forward Scatter (FSC) and Side Scatter (SSC). **b and g**. Fluorescence intensities of single bead samples FITC1-FITC5 containing beads with an increasing number of conjugated FITC fluorophores. **c and h**. Calibration curves created using MFI values of FITC1-FITC5 peaks shown in **b and g** and provided MEF values for each sample FITC1-FITC5. **d and i**. Fluorescence intensities of DNA beads with an increasing number of conjugated FAM fluorophores. **e and g**. Calibration of FCMs based on FAM calibration DNA beads shown in **d and i**.

Spherotech calibration particle detection was triggered by SSC-H (Quanteon) and FSC-H (FACSAria III), and single beads were gated in an FSC/SSC dot plot (Figure 1a and f). Spherotech FITC calibration particles were provided in 5 vials, where one vial contained blank unlabeled beads, while the beads in the other 4 vials were labeled with an increasing number of FITC fluorophores. Each vial was analyzed separately (Figure 1b and g) and a calibration curve (Figure 1c and h) was plotted based on values of molecules equivalent FITC (MEF), stated on the datasheet, and the recorded MFI values (Figure 1b and g). Spherotech “blank beads” were assigned an MEF value possibly due to blank bead autofluorescence (Figure 1b and g). FAM DNA beads with 160 Cy5 in the trigger module were analyzed as described previously in this article (Figure 1d and i). FAM DNA beads acquired on the FACSAria III were gated additionally to exclude background noise (Figure 1i). A calibration curve was plotted based on the number of FAM fluorophores on the bead and the recorded MFI values (Figure 1e and j). Calibration of the Quanteon and the FACSAria III using FITC Spherotech and FAM DNA calibration beads resulted in calibration curves with high linearity R^2^>0.997.

The biggest difference between DNA and MESF beads calibration curves was observed in the y-axis intersection: “158” vs “-21,958” for the Quanteon and “102” vs “−1,782” for the FACSAria III for calibration based on DNA beads and Spherotech particles, respectively (Figure 1). According to the calibration curve based on Spherotech FITC calibration beads, an MFI of a DNA bead with 0 FAM dyes corresponds to 12,427 MEF. The same recalculations for the FACSAria III result in 11,644 MEF per DNA bead not labeled with any FAM molecules. The slope of the DNA bead calibration curve is steeper compared to Spherotech beads suggesting that Spherotech beads may underestimate the fluorescence signal contribution per fluorophore molecule approximately 3-6 times. This can partly be explained by the contribution of trimers and tetramers these make up to 30% of all detected beads on the Quanteon and up to 20% on the FACSAria III, thus artificially increasing the steepness of the slope.

A similar calibration of the Quanteon and the Cytoflex was performed using Spherotech APC particles and Cy5 DNA calibration beads (Figure 2). Spherotech APC particles were delivered in two vials; one containing blank particles and the other containing a mix of particles labeled with 5 different APC intensities. APC single particles were gated in an FSC/SSC plot (Figure 2a and f) and MFI values of each of the 6 peaks (Figure 2b and g) were used for a calibration curve plot (Figure 2c and h). DNA beads labeled with 150 FAM in the trigger module and varying numbers of Cy5 molecules (0-70 Cy5) in the calibration module were analyzed as described earlier in this paper. The number of Cy5 fluorophores conjugated to the Cy5 DNA beads was plotted against the MFI values of these beads, resulting in a calibration curve with a high degree of linearity R^2^>0.988. As two different fluorophores were used for calibration, the comparison of Cy5 and APC calibration curves’ slope is irrelevant, but the y-axis intersection should be comparable. However, this was not the case: according to Spherotech particle calibration of the Quanteon, a DNA bead with no Cy5 dyes had a fluorescence of 10,828 molecules equivalent APC (MEAP). The same recalculations of 0 Cy5 DNA beads analyzed on the Cytoflex resulted in 10,779 MEAP.

**Figure 2.**
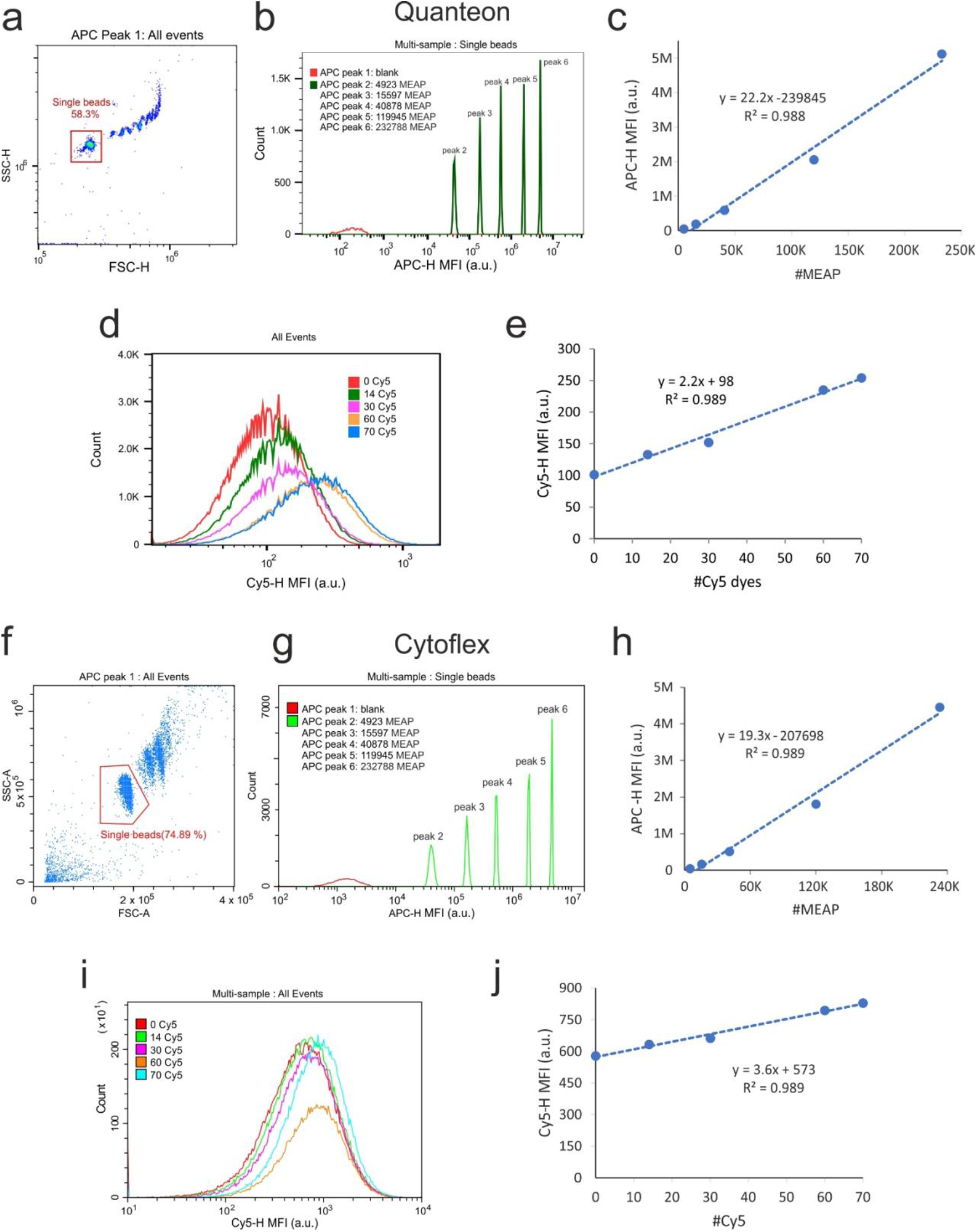
Calibration of the Quanteon (**a-e**) and the Cytoflex (**f-j**) using APC Spherotech and Cy5 DNA calibration beads. **a and f** Spherotech single APC beads were gated using Forward Scatter (FSC) and Side Scatter (SSC). **b and g** Fluorescence intensities of single peaks from the two samples. APC peak 1 and APC peak 2-6 containing beads with an increasing number of conjugated APC fluorophores. **c and h** Calibration curves were created using MFI values of APC2 –APC6 peaks shown in **b and g** and provided MEF values for each peak. **d and i** Fluorescence intensities of DNA beads with increasing number of conjugated Cy5 fluorophores. **e and g** Calibration curves created using Cy5 MFI values shown in **d and i** and a number of conjugated Cy5 fluorophores to DNA beads.

Furthermore, we have calculated Q values using the quadratic model for Q and B values[8] based on DNA and Spherotech beads calibration data shown in Figure 1 and 2 (Table 1 and 2). The Q values for the blue channel based on calculations using Spherotech data suggest that the Quanteon is approximately 4 times more sensitive than the FASCAriaIII while calculations based on DNA beads suggest that the FACSAriaIII is almost 4 times more sensitive than the Quanteon (Table 2). These disparate results may be due to the differences in the emission curves of the dyes in the Spherotech beads compared to those of the DNA beads. Our experimental data supports the calculations based on Spherotech as the Quanteon was more effective in detecting the FAM-labeled particles compared to the FACSAriaIII (Figure S8). The Q values calculated for the red channel, based on both Spherotech and DNA beads data, suggest that the Quanteon is approximately 4 times more sensitive than the Cytoflex in the Cy5 channel. That is also evident from our trigger sensitivity data for the Cy5 channel where the Quanteon is more effective in the detection of Cy5-labeled particles (Figure S9).

**Table 1.** Q values calculations using Spherotech beads and DNA beads.

|  | **Spe**<br>**(QSpe)** | **BSpe** | **BIU** | **CVi A0** | **QERF** | **BERF** | **Goodness**<br>**of Fit** | **Model Used** |
| --- | --- | --- | --- | --- | --- | --- | --- | --- |
| *Spherotech beads* |  |  |  |  |  |  |  |  |
| Aria FITC-H, Mean | 3,47E-03 | -6,91E-03 | -1,99E+00 | 3,96E-03 | 4,87E-04 | -2,79E-01 | 1,00E+00 | Quad |
| Quanteon FITC-H, Mean | 1,41E-03 | 7,47E-02 | 5,31E+01 | 5,45E-03 | 2,05E-03 | 7,74E+01 | 1,00E+00 | Quad |
| CytoFlex APC-H, Mean | 1,33E-03 | -1,09E-01 | -8,20E+01 | 1,27E-04 | 1,65E-02 | -1,02E+03 | 9,97E-01 | Linear |
| Quanteon APC-H, Mean | 4,58E-03 | -6,12E-01 | -1,33E+02 | 3,02E-04 | 6,56E-02 | -1,91E+03 | 1,00E+00 | Quad |
| Aria Median, FAM-H | 1,16E-02 | -3,05E-03 | -2,64E-01 | 6,01E-01 | 7,11E-02 | -1,62E+00 | 9,68E-01 | Linear |
| Quanteon Median, FAM-H | 1,06E-03 | -2,14E-04 | -2,02E-01 | 2,41E+00 | 1,78E-02 | -3,41E+00 | 9,54E-01 | Linear |
| CytoFLEX Median, CY-5-H | 5,87E-04 | -5,82E-05 | -9,91E-02 | -9,58E-01 | 2,65E-02 | -4,47E+00 | 9,98E-01 | Linear |
| Quanteon Median, Cy5-H | 1,02E-02 | -4,37E-03 | -4,28E-01 | 3,61E-01 | 1,04E-01 | -4,35E+00 | 9,80E-01 | Linear |

**Table 2.** Q values comparison deducted using Spherotech beads vs DNA beads.

|  | Q Spherotech | Q DNA | Q DNA /Q Spherotech |
| --- | --- | --- | --- |
| Aria FITC/FAM | 0,0005 | 0,0711 | 146,0150 |
| Quanteon FITC/FAM | 0,0021 | 0,0178 | 8,6967 |
| Cytoflex APC/Cy5 | 0,0165 | 0,0265 | 1,6075 |
| Quanteon APC/Cy5 | 0,0656 | 0,1040 | 1,5851 |
|  | Q Spherotech | Q DNA |  |
| Quanteon/Aria (FITC/FAM) | 4,21 | 0,25 |  |
| Quanteon/Cytoflex (APC/Cy5) | 3,98 | 3,92 |  |

## Conclusion

Here, we aimed to test the usage of previously described DNA calibrator beads, which allow the calibration of FCM in absolute fluorophore numbers to compare the fluorescence sensitivity of five instruments. To our knowledge, these are the only described calibration beads that provide calibration in absolute fluorophore numbers and have no autofluorescence, allowing the determination of a true “0” fluorescence background. We have tested blue and red fluorescence channels using FAM and Cy5 fluorophores. Furthermore, all five FCMs were assessed for maximum counting speed, optimal fluorescence trigger and calibration gains, and gain amplification linearity for the two channels. Lastly, we compared the slope and y-axis intersection of the calibration curves obtained using DNA beads and commercially available Spherotech MESF beads. The tested FCMs vary in laser power and laser wavelength, in detector type and detection angle and are there expected to have varying sensitivities. Not all tested FCMs were able to detect single DNA beads and not in both fluorescence channels, thus, we ended up comparing calibration on only two FCMs for each of the fluorescence channels, where both DNA and MESF beads calibration was possible.

In the blue channel, we observed a large difference in the detected y-axis intersection when comparing MESF beads and DNA beads: according to MESF bead calibration a DNA bead with 0 FAM molecules generates a signal equal to more than 11,000 MEF. However, because DNA beads lack any significant autofluorescence in the FITC/FAM channel, they provide a true zero point and, thus, should represent the real y-axis intersection for the calibration curve. Similarly, in the red channel, we observed a large difference in the y-axis intersection when comparing MESF and DNA beads. The y-axis intersection denotes the point of “zero” fluorochrome bead signal should be the same for blank beads regardless of fluorochrome not used.

It should be remembered that MESF units are not absolute fluorophore numbers and cannot be used to directly enumerate fluorophores. Especially when dim particles are analyzed, MESF beads will substantially overestimate the actual number of fluorophores because they assign 0 dye fluorescence to a signal corresponding to more than 10,000 MESF. However, MESF calibration beads can be used to translate arbitrary units to MESF units that, in turn, can be used to observe day-to-day instrument changes and instrument sensitivity allowing comparison of results across different instruments in terms of MESF units [6–8, 16].

Because DNA beads provide calibration in absolute fluorophore numbers and lack any significant background fluorescence, they are especially suitable for the characterization of dim fluorescing particles such as viruses and extracellular vesicles. However, these calibrators have some limitations: a fluorescence trigger is needed for their detection and the number of fluorophores per DNA calibrator is limited to 220 in the present setting. Therefore, one cannot increase the trigger fluorophore array substantially without decreasing the calibration fluorophore array. This requires determination of optimal trigger gain on an instrument and limits the use of the beads to FCMs that are sensitive enough to detect DNA calibrators with a maximum of 160 trigger fluorophores. Secondly, DNA beads can only be functionalized with small organic fluorophores (like Cy3, Cy5, FITC/FAM, and the Alexa Fluor dyes) and it is, thus, not possible to use DNA beads for labeling with arrays of large molecules such as fluorescent proteins like APC and PE.

The five FCMs compared in this study perform very differently. There are several reasons for this *e*.*g*. differences in laser excitation wavelengths and power, bandpass filters, and detector type. However, the results in this study show that newer FCM models and surprisingly also some older versions can readily detect DNA calibration beads and can thus be calibrated using DNA beads in absolute fluorophore numbers. We, therefore, believe that these instruments can also be used in bio-nanoparticle analysis and biochemical characterization of dim fluorescing particles.

## Methods and Materials

### Synthesis of DNA beads

The design and synthesis of DNA calibration beads were described elsewhere [1]. Briefly, DNA beads were self-assembled in a one-pot reaction by mixing scaffold strand M13 ssDNA 8064 bases (Tilibit, Germany) with 10x excess of staple strands (5x excess for Cy5 and FAM labeled HLPC purified oligos) in 50 μL TAE/Mg^2+^ buffer (40 mM Tris-acetate, 1 mM EDTA, pH = 8.3, 12-15 mM Mg^2+^) using a temperature ramp: incubation at 65°C for 20 min, followed by a steep decrease to 50°C and slow cooling down to 40°C with a temperature decrease of 0.1°C/12 min, with a hold step at 20°C before purification using MicroSpin S-400 HR (GE Healthcare Life Sciences, USA). The standard MicroSpin protocol was followed using the annealing buffer (TAE/12-15 mM Mg^2+^ buffer) for the column wash before purification. Post-purification DNA beads were stored in the dark at −80°C.

### Spherotech MESF calibration beads

Sphero™ FITC Calibration beads (cat. no. ECFP-F1-5K) and Sphero™ APC Calibration beads (cat. no. ACP-30-SK) (Spherotech, Lake Forest, IL, USA) were vigorously vortex mixed before dilution in TAE buffer.

### Flow cytometers and data analysis software

Data acquired on the NovoCyte 3000 or NovoCyte Quanteon 4025 (both ACEA Biosciences, Inc. A part of Agilent) was analyzed in NovoExpress software version 1.3.0 (ACEA Biosciences, Inc. A part of Agilent) and FlowJo (v. 10.8.1, BD). Data acquired on an LSRFortessa with HTS loader and FACSAria III (both BD Biosciences) was acquired in BD FACSDiva software version 8.0.2 (BD Biosciences) and analyzed in FlowJo (v. 10.8.1, BD). Data acquired on a Cytoflex (Beckman Coulter) was analyzed using Cytexpert 2.03.0.84 software version.

| Laser |  | Novocyte 3000 | Quanteon 4025 | LSRFortessa | FACSARIA III | CytoFlex |
| --- | --- | --- | --- | --- | --- | --- |
| 488 nm | Laser power | 20mW | 100mW | 50mW | 20mW | 50mW |
|  | Detection filter | 530/30 | 530/30 | 530/30 | 530/30 | 525/40 |
| 633-640 nm | Wavelength / power | 640nm / 40mW | 637nm / 100mW | 640nm / 40mW | 633nm / 20mW | 638nm / 50mW |
|  | Detection filter | 675/30 | 675/30 | 730/45 | 730/45 | 660/20 |

### Choice of the threshold value

For all flow cytometers, the trigger was set on either the FAM or Cy5 parameter, depending on the experiment, to achieve an event rate between 10 and 20 events/second on a buffer sample run at the lowest flow speed. For the calibration experiments, the threshold was set at 100 events/second on a buffer sample.

### FCM count rate test

Two sets of DNA beads with either 10 or 100 FAM dyes in the calibration module and 100 Cy5 dyes in the trigger module were used to assess the maximum speed of different flow cytometers. DNA beads with 0 FAM dyes and 100 Cy5 dyes were used to determine total background fluorescence. Serial dilutions of each sample were analyzed for 90 seconds.

The flow rates for each flow cytometer were set to the lowest in all experiments described in this paper: LRSFortessa 60 µl/min, Quanteon 5 µl/min, FACSAria III 5 µl/min, Novocyte 5 µl/min, and Cytoflex 10 µl/min.

### Optimal FAM trigger gain determination

Cy5 beads with 100 FAM dyes in the trigger module and a varying number of Cy5 dyes (0, 10, 40, 70, or 110) were used to determine optimal trigger gain. The threshold was set on the FAM parameter for each FAM gain, as described above, and samples were acquired for 60 seconds. Cy5 calibration gain was held constant (LSRFortessa at 700, Quanteon 1,000, FACS Aria III 850, Novocyte 800, and Cytoflex 2,000) while the FAM trigger gain was varied.

### Optimal Cy5 trigger gain determination

Optimal Cy5 trigger gain was determined in a similar way using FAM beads labeled with a varying number of FAM dyes (0, 10, 40, 70, or 100) in the calibration module and 100 Cy5 in the trigger module. The threshold was set on Cy5 for each Cy5 gain. FAM calibration gain was held constant (LSRFortessa at 700, Quanteon 1,000, FACS Aria III 850, Novocyte 800, and Cytoflex 2,000).

### FAM detector gain amplification

FAM beads with a varying number of FAM dyes (0, 10, 40, 70, or 100) and a constant number of 100 Cy5 dyes in the trigger module were used to assess FAM detector gain amplification. Cy5 trigger gain was held constant (LSRFortessa at 575, Quanteon 500, FACS Aria III 730, Novocyte 750, and Cytoflex 400) while varying FAM calibration gain.

### Cy5 detector gain amplification

Cy5 beads with a varying number of Cy5 dyes (0, 10, 40, 70, or 110) and a constant number of 100 FAM dyes in the trigger module were used to assess Cy5 detector gain amplification. FAM trigger gain was held constant (LSRFortessa at 630, Quanteon 800, FACS Aria III 700, Novocyte 600, and Cytoflex 400) while varying FAM calibration gain.

### DNA beads ensemble fluorescence measurement

DNA beads, diluted using TAE buffer spiked with 13mM Mg^2+^ and 60 µL of beads (0.01-0.05 nM), were loaded into a quartz cuvette with a volume of 60 μL (Hellma). Fluorescence spectra profiles of both Cy5 and FAM fluorophores were recorded using a spectrofluorometer (FluoroMax-3 from HORIBA Jobin Yvon) after excitation of FAM and Cy5 at 488 nm or 600 nm, respectively.

### Agarose gel electrophoresis assay

Assembled DNA beads were loaded onto SybrSafe (Invitrogen) prestained 1% agarose gel containing 5 mM Mg^2+^ and allowed to migrate for 3.5 h, 4°C at 80 V. The gel was imaged using ChemiDoc XRS+ (Bio-Rad laboratories) and analyzed using ImageLab 6.0.1 software version (Bio-Rad laboratories)

## Notes

### Competing Interest Statement

The authors have declared no competing interest.

