## Supplementary for "Inter-Instrument Quantification of Fluorescence Read-out Signal Using DNA Origami Calibrator Beads"

Supplementary Material

Bohn A.B., Petersen C.C., Wood J., Pedersen F.S., Selnihhin D.

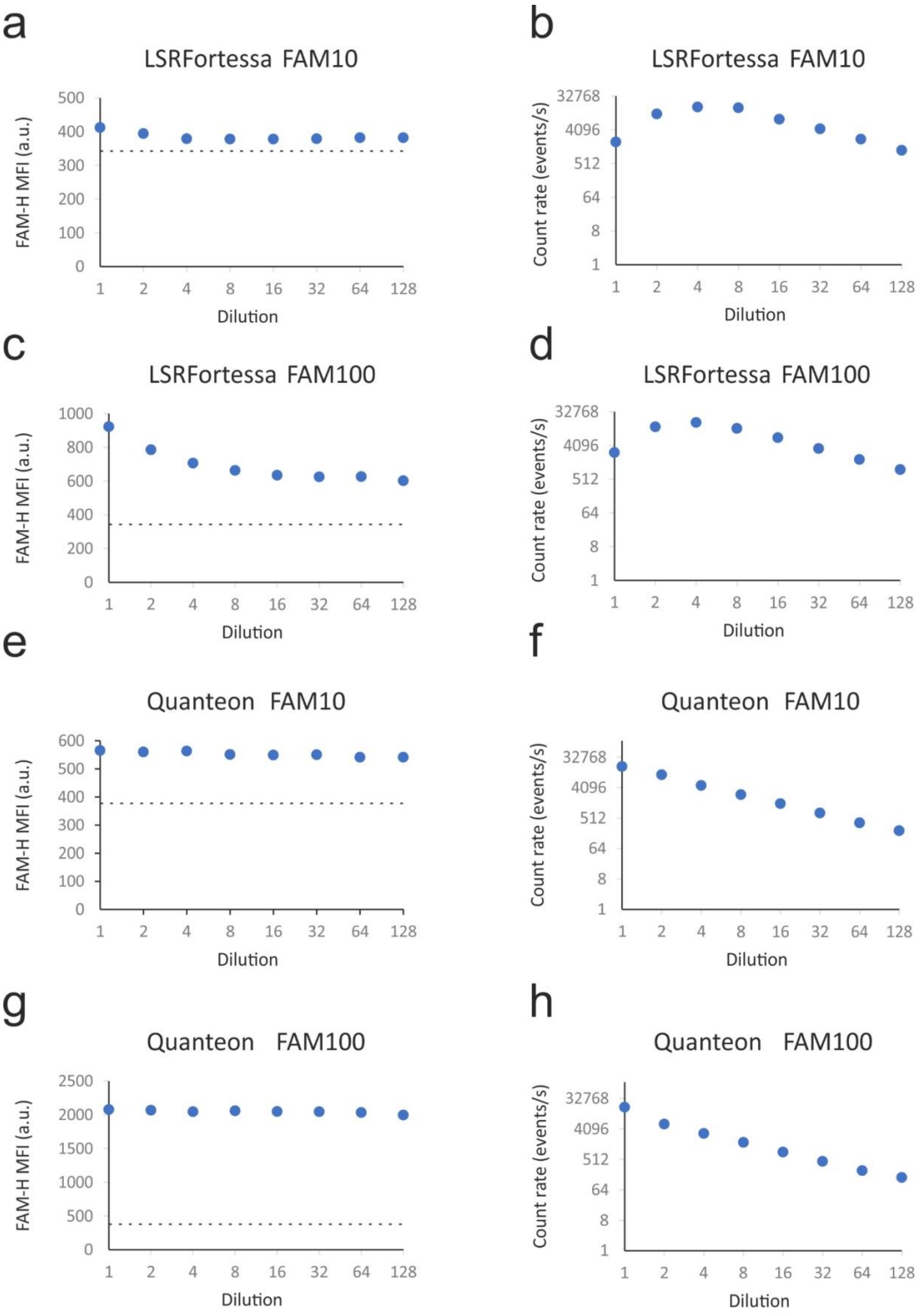

a
LSRFortessa FAM10
FAM-H MFI (a.u.)
0
100
200
300
400
500
Dilution
b
LSRFortessa FAM10
Count rate (events/s)
1
8
64
512
4096
32768
Dilution
c
LSRFortessa FAM100
FAM-H MFI (a.u.)
0
200
400
600
800
1000
Dilution
d
LSRFortessa FAM100
Count rate (events/s)
Dilution
e
Quanteon FAM10
FAM-H MFI (a.u.)
0
100
200
300
400
500
600
Dilution
f
Quanteon FAM10
Count rate (events/s)
Dilution
g
Quanteon FAM100
FAM-H MFI (a.u.)
0
500
1000
1500
2000
2500
Dilution
h
Quanteon FAM100
Count rate (events/s)
Dilution
1
2
4
8
16
32
64
128

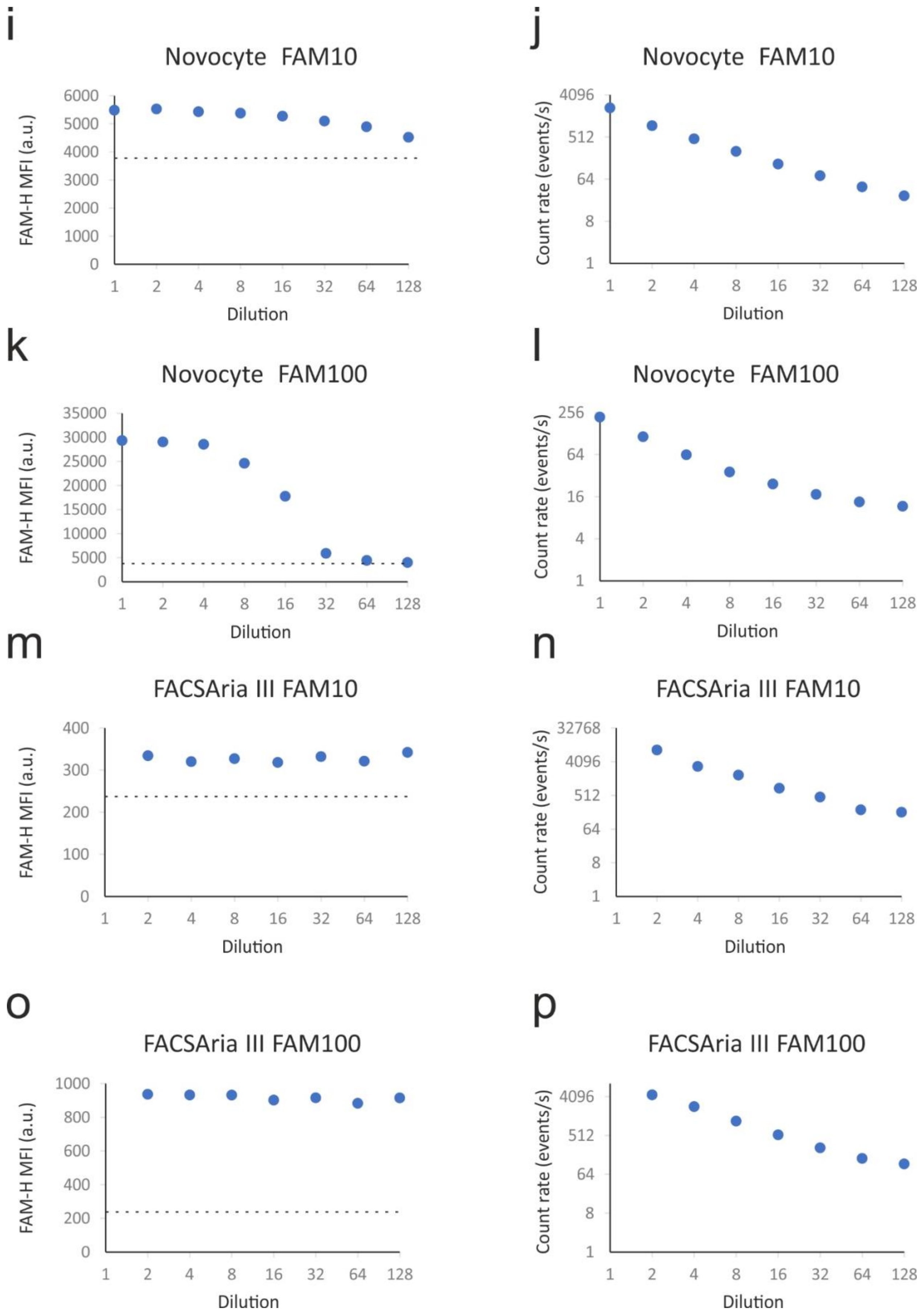
i
Novocyte FAM10
FAM-H MFI (a.u.)
0 1000 2000 3000 4000 5000 6000
1 2 4 8 16 32 64 128
Dilution
j
Novocyte FAM10
Count rate (events/s)
1 8 64 512 4096
1 2 4 8 16 32 64 128
Dilution
k
Novocyte FAM100
FAM-H MFI (a.u.)
0 5000 10000 15000 20000 25000 30000 35000
1 2 4 8 16 32 64 128
Dilution
l
Novocyte FAM100
Count rate (events/s)
1 4 16 64 256
1 2 4 8 16 32 64 128
Dilution
m
FACSAria III FAM10
FAM-H MFI (a.u.)
0 100 200 300 400
1 2 4 8 16 32 64 128
Dilution
n
FACSAria III FAM10
Count rate (events/s)
1 8 64 512 4096 32768
1 2 4 8 16 32 64 128
Dilution
o
FACSAria III FAM100
FAM-H MFI (a.u.)
0 200 400 600 800 1000
1 2 4 8 16 32 64 128
Dilution
p
FACSAria III FAM100
Count rate (events/s)
1 8 64 512 4096
1 2 4 8 16 32 64 128
Dilution

**Figure S1**. Flow cytometer (FCM) detection speed test. Two sets of DNA beads with either 10 or 100 FAM dyes in the calibration module and 100 Cy5 dyes in the trigger module were used to evaluate the highest detection speed for each FCM setup. Particle detection was triggered by Cy5 fluorescence. Serial dilutions of each sample were prepared, and each dilution was analyzed for 90 seconds. Beads with 0 FAM and 100 Cy5 dyes were used to determine the background fluorescence shown as a black dashed line in the graphs. Two sets of data: Dilution vs FAM-H Median Fluorescence Intensity (MFI) (left panels) and Dilution vs Count rate (right panels) are plotted.

All tested FCMs (Cytoflex data published previously [1]) were able to run up to 6,000 event/s without detector swarming effects except the Novocyte where approx. 2,000 events/s was the maximum recorded count rate.

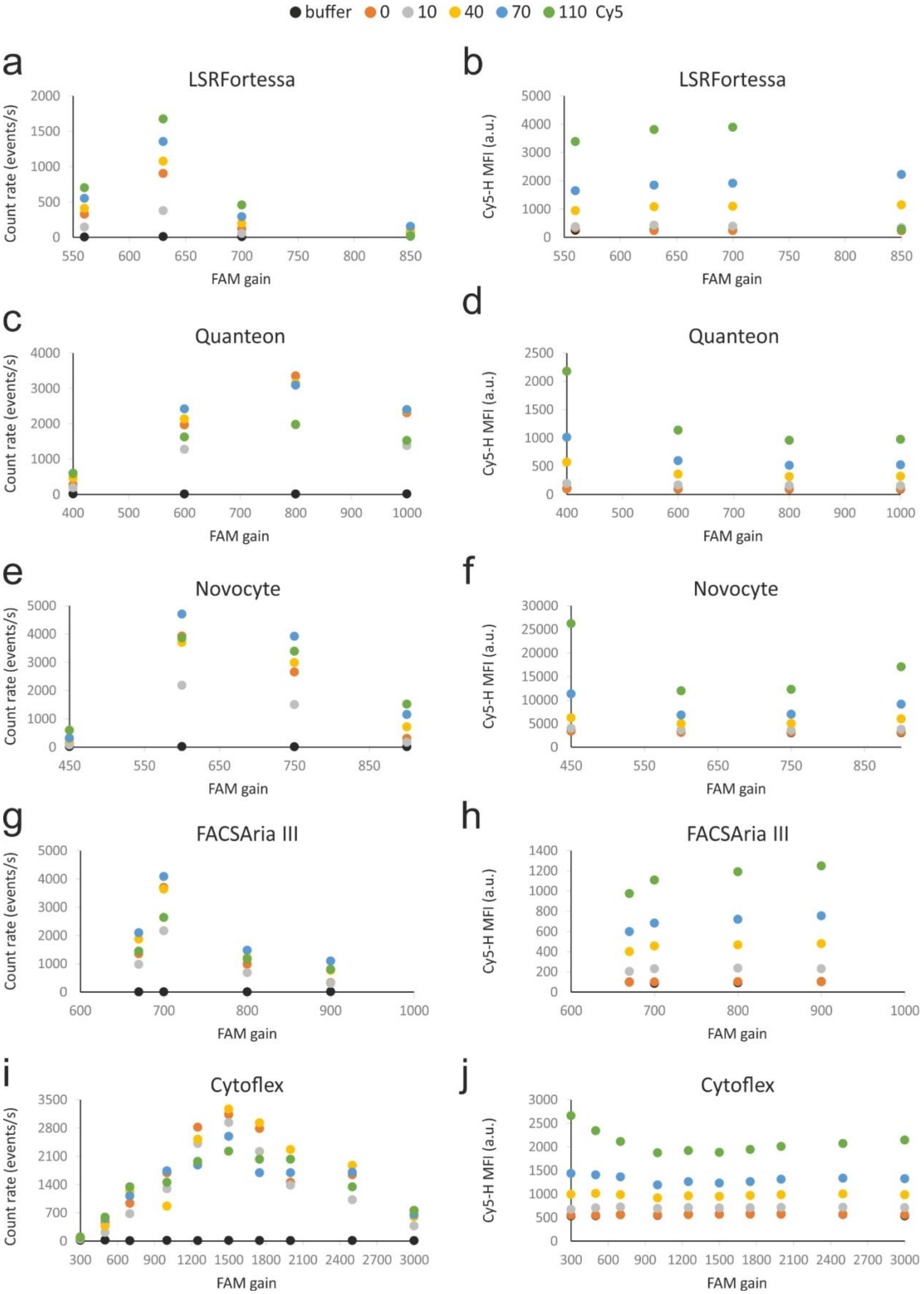

buffer
0
10
40
70
110 Cy5
a
LSRFortessa
Count rate (events/s)
FAM gain
b
LSRFortessa
Cy5-H MFI (a.u.)
FAM gain
c
Quanteon
Count rate (events/s)
FAM gain
d
Quanteon
Cy5-H MFI (a.u.)
FAM gain
e
Novocyte
Count rate (events/s)
FAM gain
f
Novocyte
Cy5-H MFI (a.u.)
FAM gain
g
FACSAria III
Count rate (events/s)
FAM gain
h
FACSAria III
Cy5-H MFI (a.u.)
FAM gain
i
Cytoflex
Count rate (events/s)
FAM gain
j
Cytoflex
Cy5-H MFI (a.u.)
FAM gain

**Figure S2**. Determination of optimal FAM trigger gain. Several FAM gains were tested to determine the most optimal gain for the detection of DNA beads with a FAM fluorescence trigger module. For each FAM gain, a threshold was set using buffer alone corresponding to the count rate of approximately 20 events/s (•). Beads with 100 FAM dyes in the trigger module and a varying number of Cy5 dyes (•0, •10, •40, •70, and •110 Cy5) in the calibration module were run at different FAM gains and constant Cy5 gain. Two data sets are plotted for each FCM: FAM trigger gain vs count rate and FAM trigger gain vs Cy5 MFI.

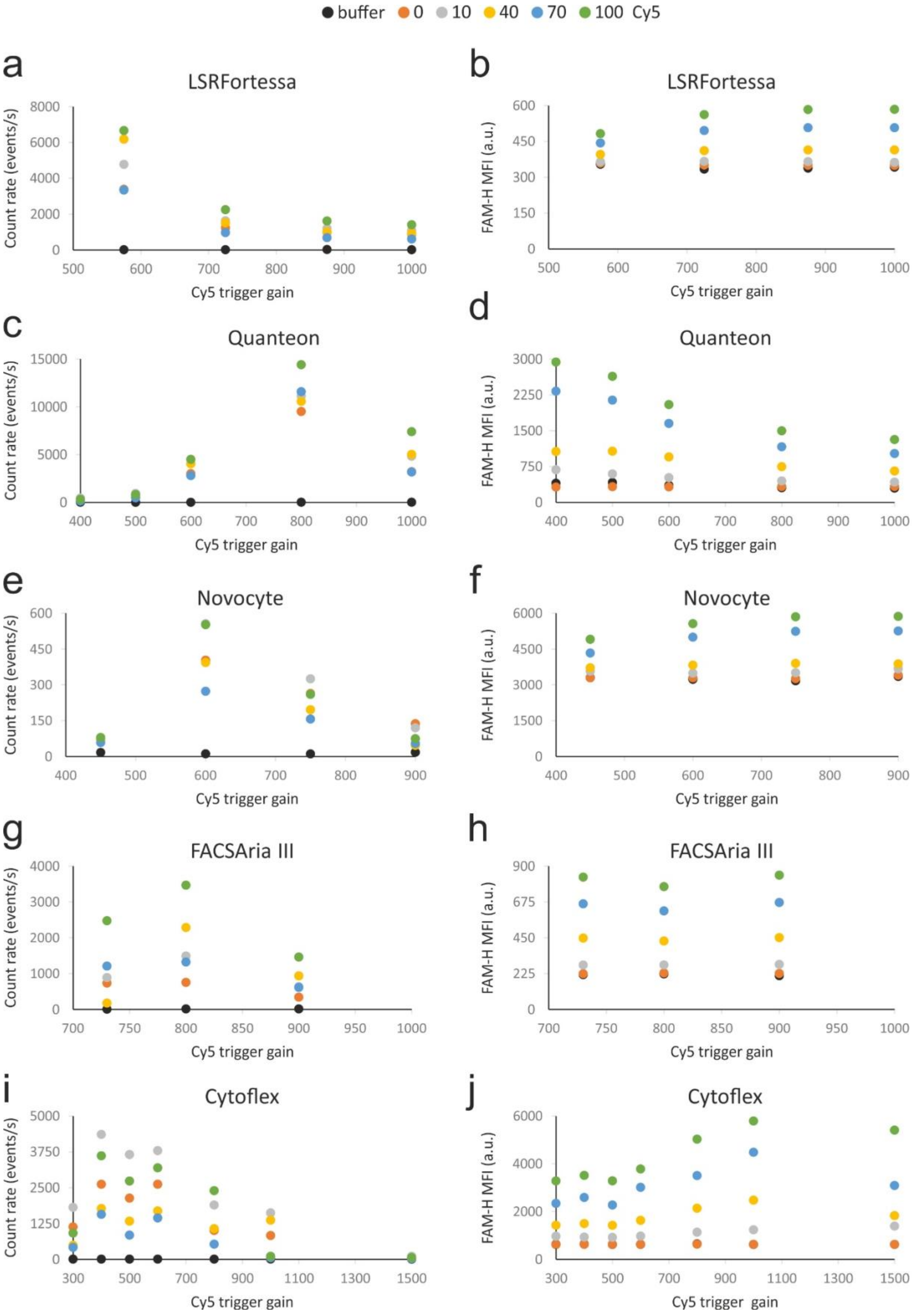
buffer 0 10 40 70 100 Cy5
a
LSRFortessa
Count rate (events/s)
Cy5 trigger gain
b
LSRFortessa
FAM-H MFI (a.u.)
Cy5 trigger gain
c
Quanteon
Count rate (events/s)
Cy5 trigger gain
d
Quanteon
FAM-H MFI (a.u.)
Cy5 trigger gain
e
Novocyte
Count rate (events/s)
Cy5 trigger gain
f
Novocyte
FAM-H MFI (a.u.)
Cy5 trigger gain
g
FACSAria III
Count rate (events/s)
Cy5 trigger gain
h
FACSAria III
FAM-H MFI (a.u.)
Cy5 trigger gain
i
Cytoflex
Count rate (events/s)
Cy5 trigger gain
j
Cytoflex
FAM-H MFI (a.u.)
Cy5 trigger gain

**Figure S3**. Determination of optimal Cy5 trigger gain. We have tested several Cy5 gains to determine the most optimal gain for the detection of DNA beads with a Cy5 fluorescence trigger module. For each gain, a threshold was set using buffer alone corresponding to a count rate of approximately 20 events/s (•). FAM beads with a varying number of FAM dyes (•0, •10, •40, •70, and •100 FAM) and 100 Cy5 dyes in the trigger module were run at different Cy5 gains and a constant FAM gain. Two data sets are plotted for each FCM: Cy5 trigger gain vs Count rate and Cy5 trigger gain vs FAM MFI.

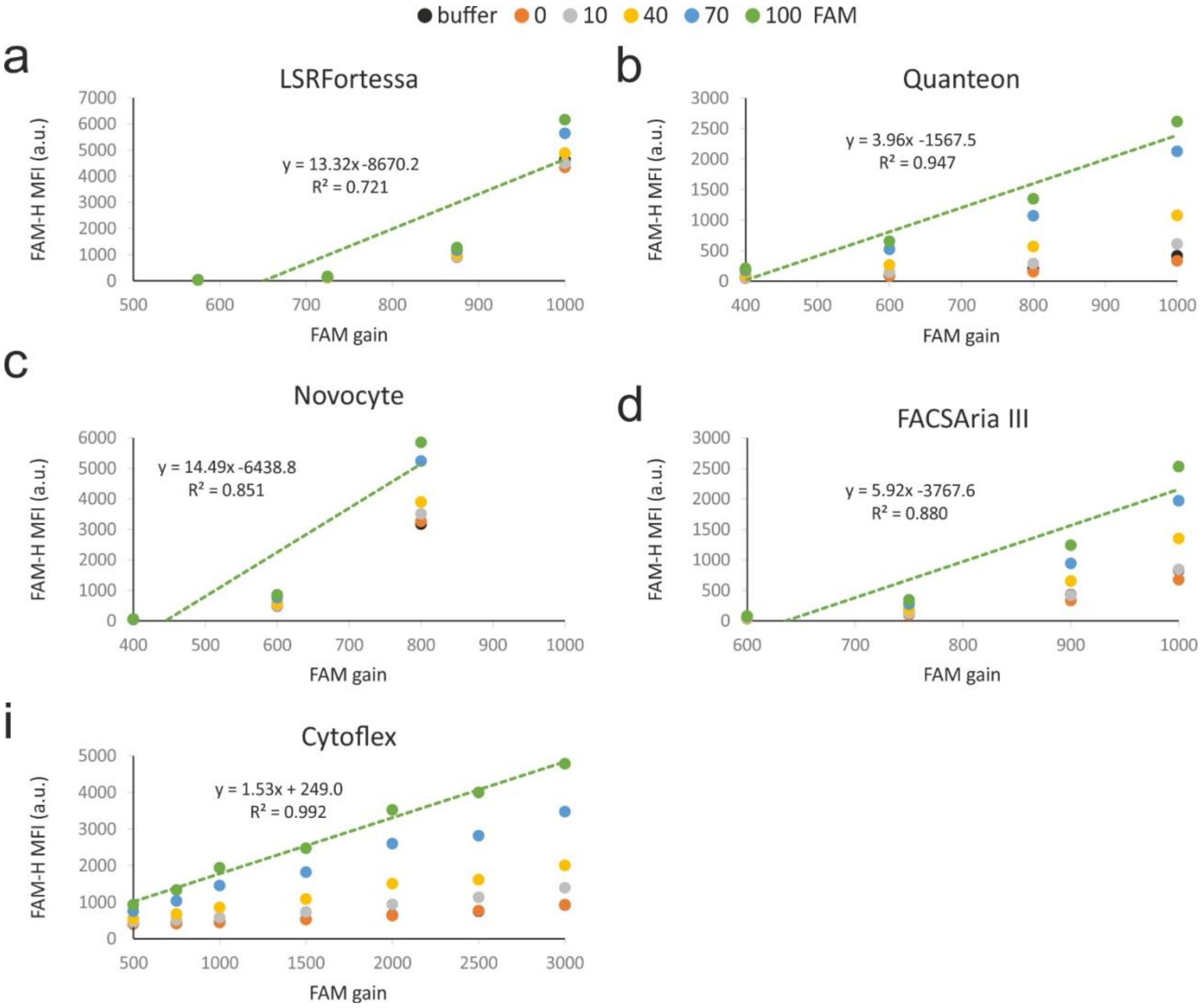


**Figure S4**. FAM detector gain amplification. To test FAM detector gain amplification, FAM beads with varying number of FAM fluorophores (•0, •10, •40, •70, and •100 FAM) and 100 Cy5 dyes in the trigger module were run at different FAM gains and constant Cy5 trigger gain. The Green dashed line represents a trend line with an accompanying equation and $R^2$ values.

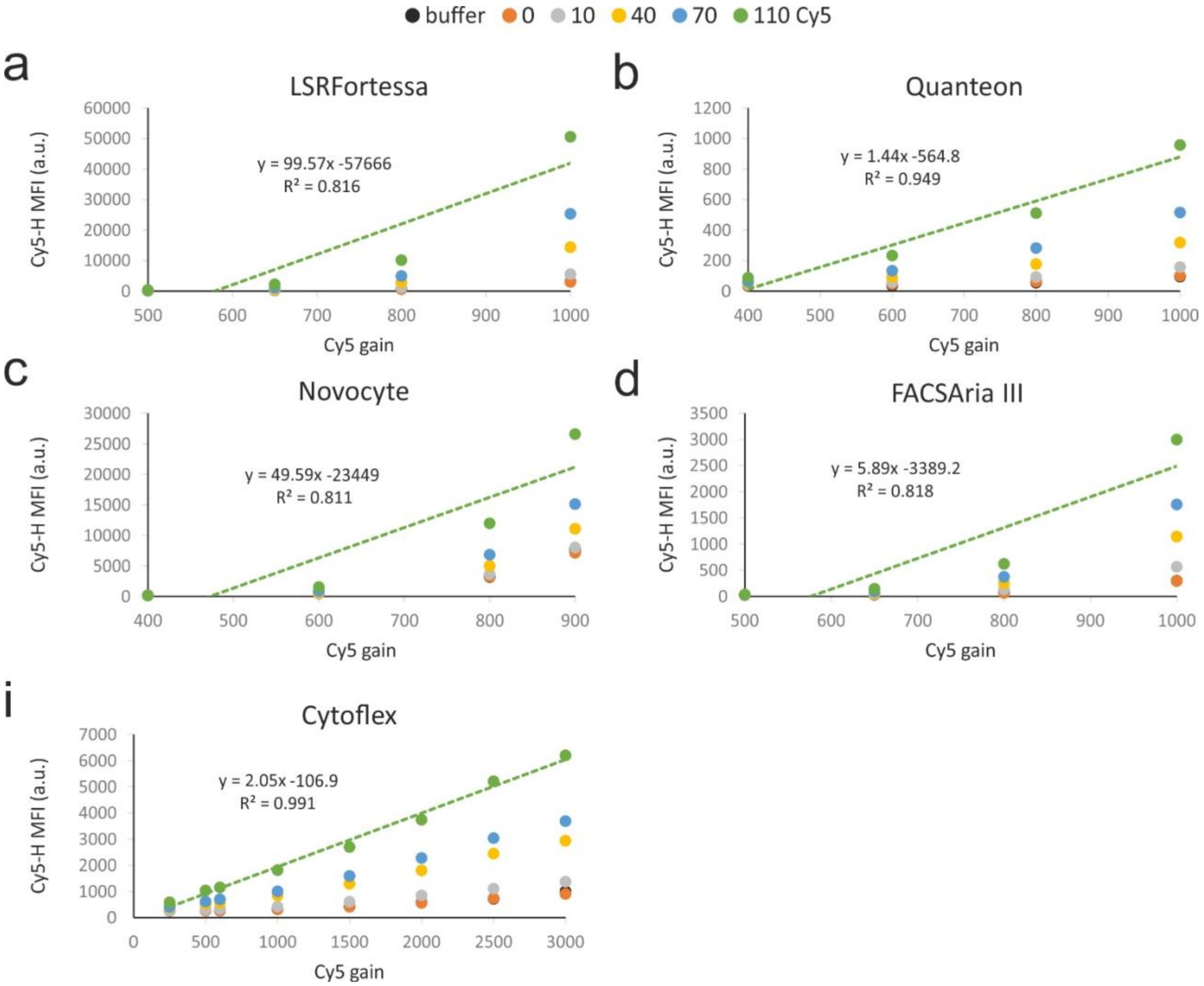


**Figure S5**. Cy5 detector gain amplification. To test Cy5 detector gain amplification Cy5 beads with varying number of Cy5 fluorophores (•0, •10, •40, •70, and •110 Cy5) and 100 FAM dyes in the trigger module were run at different Cy5 gains and constant FAM trigger gain.

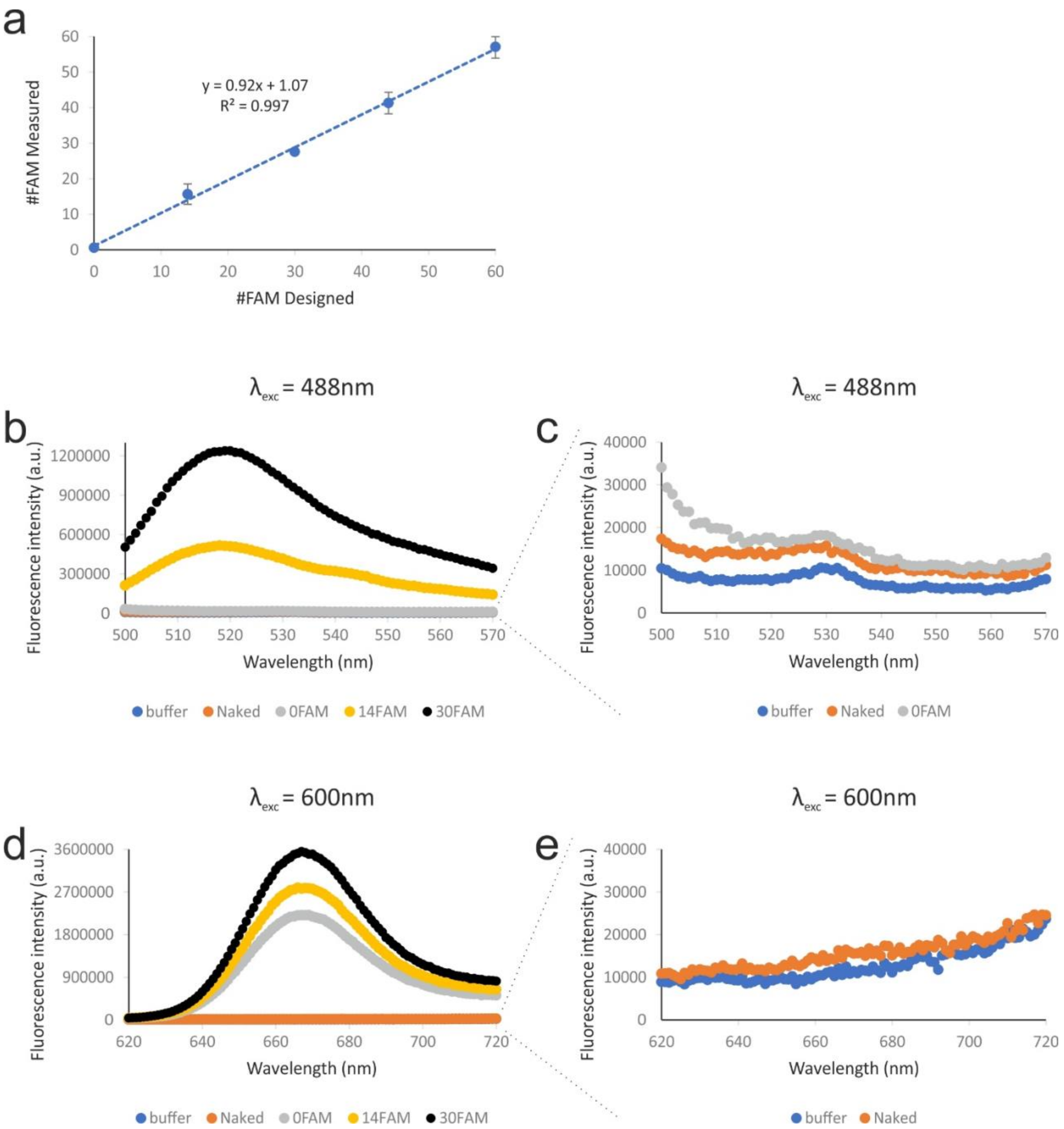

a
y = 0.92x + 1.07
$R^2$ = 0.997
#FAM Measured
#FAM Designed
λexc = 488nm
b
Fluorescence intensity (a.u.)
Wavelength (nm)
buffer Naked 0FAM 14FAM 30FAM
λexc = 488nm
c
Fluorescence intensity (a.u.)
Wavelength (nm)
buffer Naked 0FAM
λexc = 600nm
d
Fluorescence intensity (a.u.)
Wavelength (nm)
buffer Naked 0FAM 14FAM 30FAM
λexc = 600nm
e
Fluorescence intensity (a.u.)
Wavelength (nm)
buffer Naked

**Figure S6**. Ensemble characterization of FAM calibration beads and their autofluorescence. Bulk fluorescence measurements of FAM beads with 160 Cy5 in the trigger module and a varying number of FAM fluorophores were performed using a fluorimeter. **a**. "#FAM measured" was calculated as FAM/Cy5 fluorescence ratio at 520 and 670 nm, respectively, and normalized to beads containing 14 FAM fluorophores. The average of three experiments is plotted with s.d. error bars. **b**. Fluorescence intensity profiles of five samples: buffer, naked DNA particle – no dyes just pure DNA particles, DNA particles with 0 FAM and 160 Cy5 dyes, DNA particles with 14 FAM and 160 Cy5 dyes, and DNA particles with 30 FAM and 160 Cy5 excited at 488 nm. **c**. Zoom in on data from **b,** showing that beads with 0 FAM dyes and unlabeled DNA beads lack any significant fluorescence emission at FAM emission wavelength. **d**. Fluorescence intensity profiles of the same samples as in **b** excited at 600 nm. **e.** Zoom in on data from **d,** showing naked DNA beads lacking any fluorescence emission at Cy5 emission wavelength.

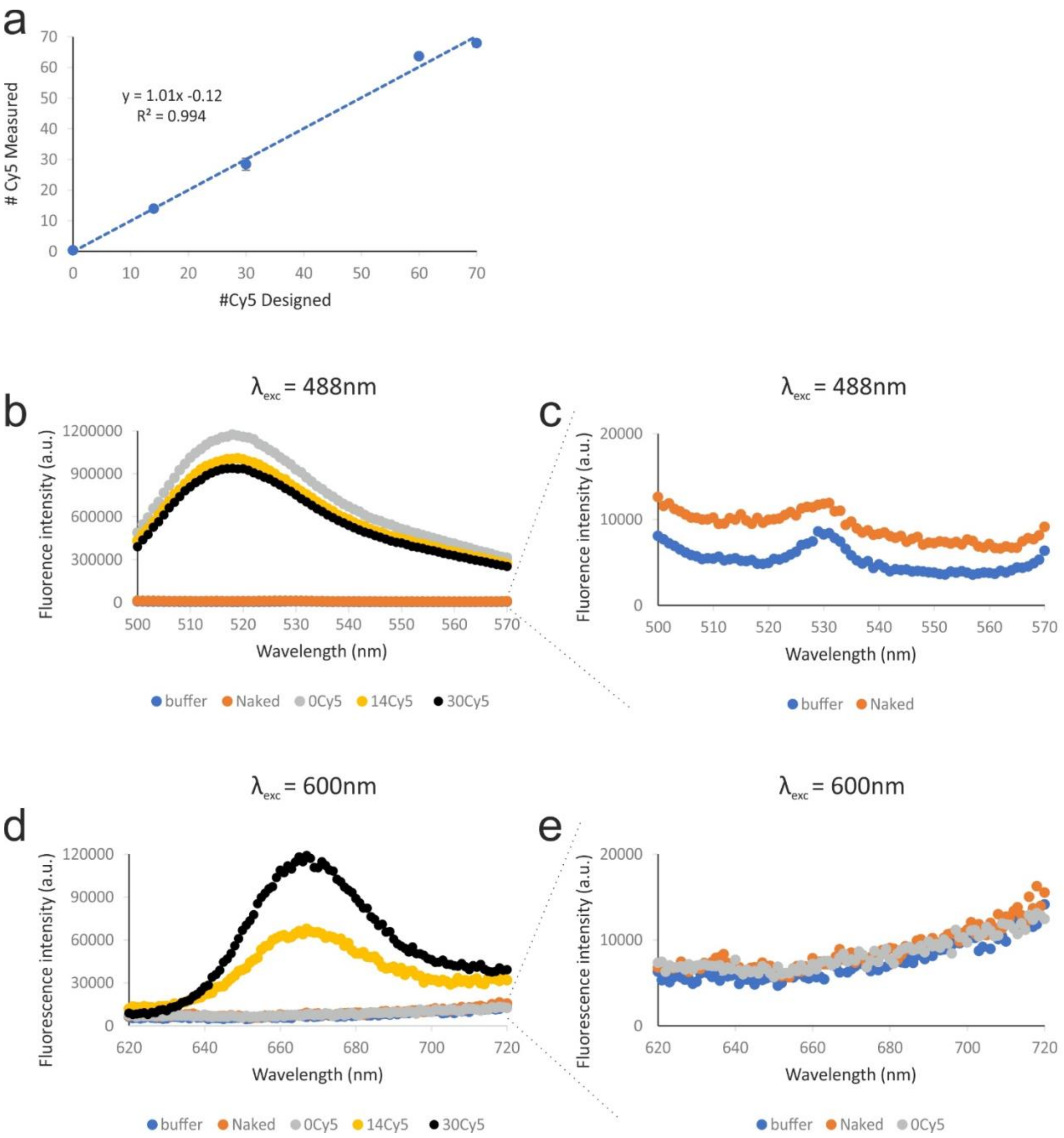

a
y = 1.01x -0.12
R² = 0.994
### Cy5 Measured
#Cy5 Designed
b
λexc = 488nm
Fluorence intensity (a.u.)
Wavelength (nm)
buffer
Naked
0Cy5
14Cy5
30Cy5
c
λexc = 488nm
Fluorence intensity (a.u.)
Wavelength (nm)
buffer
Naked
d
λexc = 600nm
Fluorescence intensity (a.u.)
Wavelength (nm)
buffer
Naked
0Cy5
14Cy5
30Cy5
e
λexc = 600nm
Fluorescence intensity (a.u.)
Wavelength (nm)
buffer
Naked
0Cy5

**Figure S7**. Ensemble characterization of Cy5 calibration beads and their autofluorescence. Bulk fluorescence measurements of Cy5 beads with 150 FAM in the trigger module and a varying number of Cy5 fluorophores were performed using a fluorimeter. **a**. "#Cy5 measured" was calculated as Cy5/FAM fluorescence ratio at 670 and 520 nm, respectively, and normalized to beads containing 14 Cy5 dyes. The average of three experiments is plotted with s.d. error bars. **b**. Fluorescence intensity profiles of five samples: buffer, naked DNA particles, DNA particles with 0 Cy5 and 150 FAM dyes, DNA particles with 14 Cy5 and 150 FAM dyes, and DNA particles with 30 Cy5 and 150 FAM excited at 488 nm. **c**. Zoom in on data from **b,** showing that unlabeled DNA beads lack any significant fluorescence emission at FAM emission wavelength. **d**. Fluorescence intensity profiles of the same samples as in **b** excited at 600 nm. **e.** Zoom in on data from **d,** showing unlabeled DNA beads and beads with 0 Cy5 lacking any fluorescence emission at Cy5 wavelength.

a

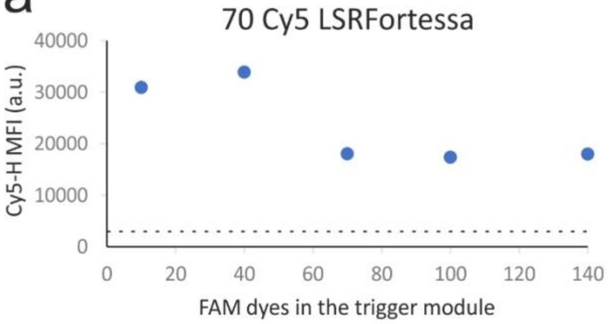


| Detected concentration | | |
|---|---|---|
| | pM | % |
| 10 FAM | 0.005 | 0.002 |
| 40 FAM | 0.26 | 0.1 |
| 70 FAM | 0.60 | 0.3 |
| 100 FAM | 0.98 | 0.5 |
| 140 FAM | 2.79 | 1.3 |

b

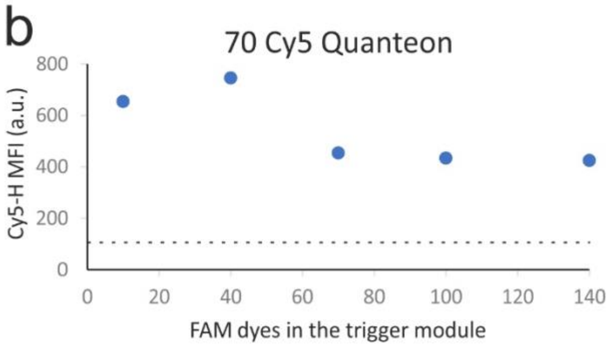


| Detected concentration | | |
|---|---|---|
| | pM | % |
| 10 FAM | 0.26 | 0.1 |
| 40 FAM | 17.21 | 8.0 |
| 70 FAM | 13.48 | 6.3 |
| 100 FAM | 72.37 | 33.7 |
| 140 FAM | 94.76 | 44.1 |

c

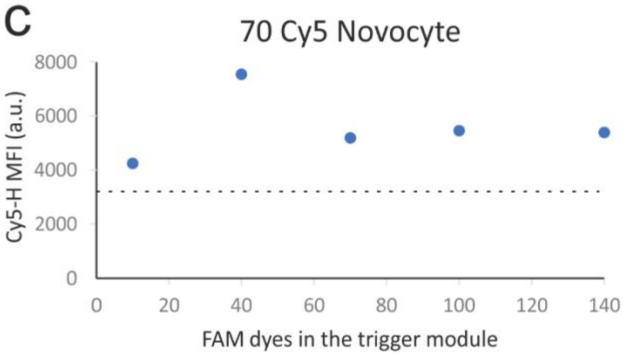


| Detected concentration | | |
|---|---|---|
| | pM | % |
| 10 FAM | 0.03 | 0.01 |
| 40 FAM | 0.99 | 0.5 |
| 70 FAM | 1.80 | 0.8 |
| 100 FAM | 7.47 | 3.5 |
| 140 FAM | 16.79 | 7.3 |

d

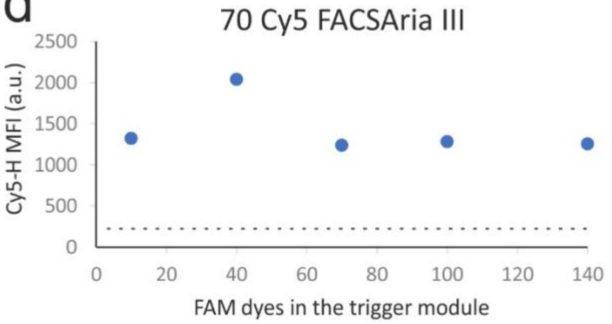


| Detected concentration | | |
|---|---|---|
| | pM | % |
| 10 FAM | 0.12 | 0.1 |
| 40 FAM | 3.34 | 1.6 |
| 70 FAM | 4.01 | 1.9 |
| 100 FAM | 6.64 | 3.1 |
| 140 FAM | 10.59 | 4.9 |

i

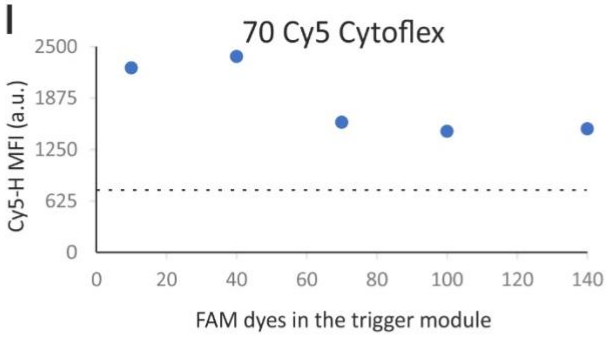


| Detected concentration | | |
|---|---|---|
| | pM | % |
| 10 FAM | 2.13 | 1.0 |
| 40 FAM | 25.92 | 12.1 |
| 70 FAM | 42.48 | 19.8 |
| 100 FAM | 74.58 | 34.7 |
| 140 FAM | 172.30 | 80.1 |

**Figure S8**. FAM trigger sensitivity. Beads with 70 Cy5 dyes and a varying number of FAM dyes (10, 40, 70, 100, and 140 FAM) in the trigger module, at a concentration of 215 pM, were used to test FAM detector sensitivity and determine the number of FAM dyes needed in the trigger module for single DNA beads to be detected. Beads with 0 Cy5 dyes and 100 FAM in the trigger module were used to determine background fluorescence illustrated with black dashed lines in the graphs.

a

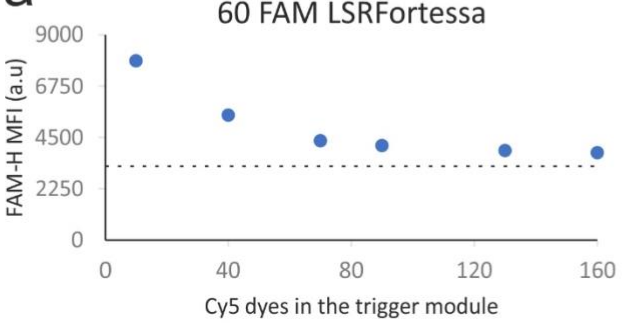


| Detected concentration | | |
|---|---|---|
| | pM | % |
| 10 Cy5 | 0.86 | 0.4 |
| 40 Cy5 | 8.61 | 4.0 |
| 70 Cy5 | 16.08 | 7.5 |
| 90 Cy5 | 27.18 | 12.6 |
| 130 Cy5 | 52.33 | 24.3 |
| 160 Cy5 | 84.59 | 39.3 |

b

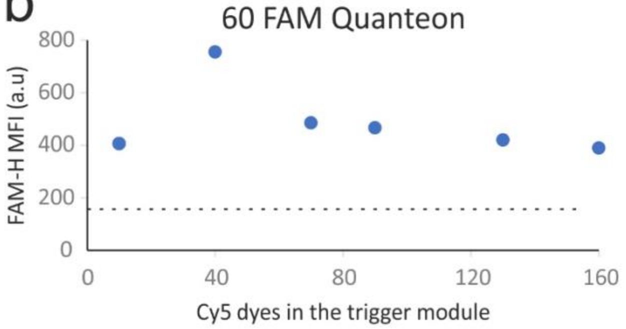


| Detected concentration | | |
|---|---|---|
| | pM | % |
| 10 Cy5 | 0.22 | 0.1 |
| 40 Cy5 | 7.47 | 3.5 |
| 70 Cy5 | 14.41 | 6.7 |
| 90 Cy5 | 28.65 | 13.3 |
| 130 Cy5 | 65.80 | 30.6 |
| 160 Cy5 | 107.16 | 49.8 |

c

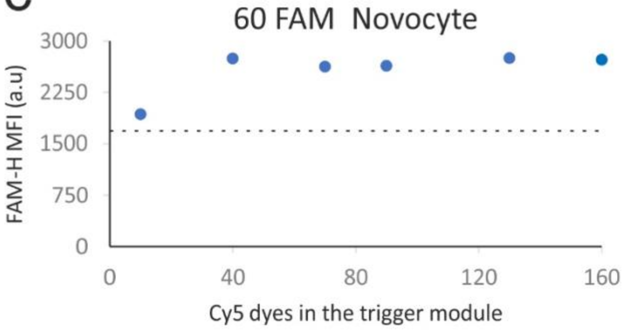


| Detected concentration | | |
|---|---|---|
| | pM | % |
| 10 Cy5 | 0.02 | 0.01 |
| 40 Cy5 | 0.28 | 0.1 |
| 70 Cy5 | 0.65 | 0.3 |
| 90 Cy5 | 1.19 | 0.6 |
| 130 Cy5 | 3.11 | 1.4 |
| 160 Cy5 | 6.02 | 2.8 |

d

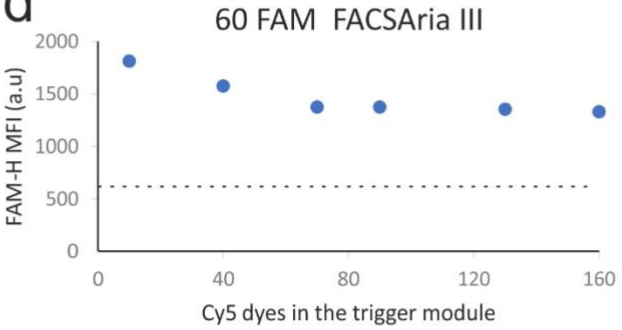


| Detected concentration | | |
|---|---|---|
| | pM | % |
| 10 Cy5 | 1.28 | 0.6 |
| 40 Cy5 | 7.38 | 3.4 |
| 70 Cy5 | 16.43 | 7.6 |
| 90 Cy5 | 35.24 | 16.4 |
| 130 Cy5 | 87.00 | 40.5 |
| 160 Cy5 | 153.12 | 71.2 |

i

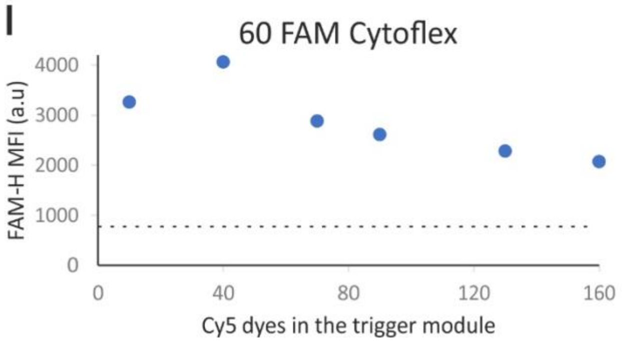


| Detected concentration | | |
|---|---|---|
| | pM | % |
| 10Cy5 | 0.79 | 0.4 |
| 40Cy5 | 4.14 | 1.9 |
| 70Cy5 | 8.27 | 3.8 |
| 90Cy5 | 8.03 | 3.7 |
| 130Cy5 | 18.14 | 8.4 |
| 160Cy5 | 24.82 | 11.5 |

**Figure S9**. Cy5 trigger sensitivity. Beads with 60 FAM and a varying number of Cy5 dyes (10, 40, 70, 90, 130, and 160 Cy5) in the trigger module, at a concentration of 215 pM, were used to determine Cy5 detector sensitivity and number of Cy5 dyes needed in the trigger module for single DNA particles to be detected. Beads with 0 FAM dyes and 100 Cy5 in the trigger module were used to determine background fluorescence as illustrated with black dashed lines in the graphs.

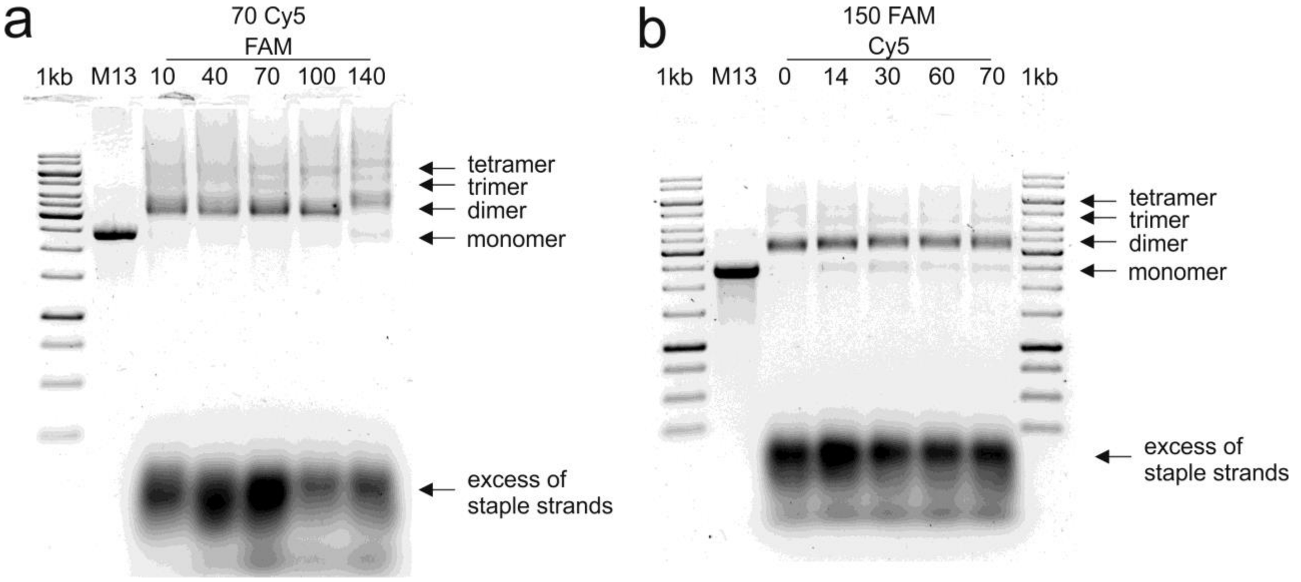


**Figure S10**. Gel electrophoresis analysis of self-assembled DNA beads. **a.** Beads with 70 Cy5 and varying numbers of FAM (10, 40, 70, 100, and 140) dyes in the trigger module, M13 –scaffold strand for DNA beads assembly. **b.** Beads with 150 FAM in the trigger module and varying numbers of Cy5 (0, 14, 30, 60, and 70) dyes in the detector module.
